# Rapid Time-stamped Analysis of Filament Motility

**DOI:** 10.1101/399006

**Authors:** Gijs Ijpma, Zsombor Balassy, Anne-Marie Lauzon

## Abstract

The in vitro motility assay is a valuable tool to understand motor protein mechanics, but existing algorithms are not optimized for accurate time resolution. We propose an algorithm that combines trace detection with a time-stamped analysis. By tracking filament ends, we minimize data loss from overlapping and crossing filaments. A movement trace formed by each filament end is created by time-stamping when the filament either first (filament tip) or last (filament tail) occupies a pixel. A frame number vs distance curve is generated from this trace, which is segmented into regions by slope to detect stop-and-go movement. We show, using generated mock motility videos, accurate detection of velocity and motile fraction changes for velocities <0.05 pixels per frame, without manual trace dropping and regardless of filament crossings. Compared with established algorithms we show greatly improved accuracy in velocity and motile fraction estimation, with greatly reduced user effort. We tested two actual motility experiments: 1) Adenosine triphosphate (ATP) added to skeletal myosin in rigor; 2) myosin light chain phosphatase (MLCP) added to phasic smooth muscle myosin. Our algorithm revealed previously undetectable features: 1) rapid increase in motile fraction paralleled by a slow increase in velocity as ATP concentration increases; 2) simultaneous reductions in velocity and motile fraction as MLCP diffuses into the motility chamber at very low velocities. Our algorithm surpasses existing algorithms in the resolution of time dependent changes in motile fraction and velocity at a wide range of filament lengths and velocities, with minimal user input and CPU time.

## Introduction

In vitro motility or gliding assays are essential tools for quantifying mechanical parameters of the interaction of motor molecules with propelled filaments. A number of (semi)-automated algorithms have been developed to track filaments as they are propelled over the motility surface (Hamelink et al. 1999; Homsher, Wang, and Sellers 1992; Månsson and Tågerud 2003; Marston et al. 1996; Mashanov and Molloy 2007; Roman et al. 2013; Root and Wang 1994; Ruhnow, Zwicker, and Diez 2011; von Wegner et al. 2008; Work and Warshaw 1992; Hilbert, Bates, et al. 2013). Most of these algorithms rely on identification of the filament in each frame by converting a grey-scale image of fluorescently labelled filaments to a black and-white image with either a manually or automatically set grey value as threshold between black and white (thresholding). Each area of connected white pixels outlines the approximate location of each filament and is individually labelled (segmentation) and correlated between frames. Some algorithms calculate the centroid of each white area and a velocity is calculated from the movement of the centroid location frame-to-frame, either with very low frame rates or considerable smoothing of the data with a moving average filter (Homsher, Wang, and Sellers 1992; Månsson and Tågerud 2003; Marston et al. 1996; Mashanov and Molloy 2007; Root and Wang 1994; Work and Warshaw 1992; Hilbert, Bates, et al. 2013). This method does not distinguish between directed (gliding) or undirected (Brownian noise, circling around a fixed point, *etc.*) motion. Furthermore, fluctuations in fluorescence intensity or small changes in microscope focus can result in a change of shape and/or size of the white area without an actual change in the filament position or size, leading to motion artifacts. Difficulties arise when filaments cross or overlap, resulting in exclusion of both filaments from the analysis. Crossing traces are either removed by hand (Homsher, Wang, and Sellers 1992; Roman et al. 2013; Work and Warshaw 1992; Hilbert, Bates, et al. 2013) or with an automated algorithm (Månsson and Tågerud 2003; Hilbert, Bates, et al. 2013). More advanced algorithms can track filaments despite crossing but require some assumptions on filament rigidity (near straight line movement) (Butt et al. 2010; Mashanov and Molloy 2007). Lastly, any curved motion results in underestimation as the centroid of a curved filament follows a shorter track than any point of the filament, as it essentially cuts corners. Some of these issues (cutting corners, lighting and focus changes) can be countered by taking a centre point on the filament instead of the centroid (Work and Warshaw 1992; Hilbert, Bates, et al. 2013).

More sophisticated software use thresholding to define a region of interest (ROI) with the approximate location of the filament, followed by identification of a more exact location of the filament by fitting segments to a model (Ruhnow, Zwicker, and Diez 2011). In theory this gives a much more accurate position estimation for the filament, it allows for analysis of crossing filaments, and it can provide detailed information about filament shape (curvature, length) and degree of directed vs. non-directed motion. However, this method is CPU intensive and it is unclear what the effect of noise is on the position accuracy. Fully automated filament detection through segmentation has been applied to single images of filament and filament like structures ((Xiao et al. 2016) among others), but the CPU intensiveness of this approach has prevented it from being useful for motility studies.

To reduce the effect of non-directed motion on calculated filament sliding velocities, one can look at the traces generated over multiple frames by each filament (Hamelink et al. 1999; Hilbert, Cumarasamy, et al. 2013; Homsher, Wang, and Sellers 1992; von Wegner et al. 2008). The trace length minus the filament length is equal to the total moved distance by the filament and is indicative of filament velocity. This averaged velocity does not distinguish between stop-and-go movement and slow movement, and it only provides a single velocity without time resolution. We previously proposed a combined algorithm using the above described centroid method to determine motile fraction for each filament, i.e. the fraction of time the velocity of movement was above a certain threshold value and combined this with the trace method to determine the velocity of the filament when moving (Hilbert, Cumarasamy, et al. 2013). However, when studying slow moving motility from actin propelled by slow motor proteins, directional motion is often less then non-directional motion on a frame-to-frame basis, particularly for small filaments. Consequently, this method does not provide an accurate estimate of velocity over time, particularly at low velocities.

Here we propose a novel approach to filament tracking which maps the motion of filament ends by storing a time stamped trace of the tip and tail of filaments. This allows for rapid analysis of actual filament velocities over time, minimal rejection from filament crossings, reduced bias from non-directional motion, and all this regardless of the curvature of the filament and filament path.

## Methods

### Motility preparation

All motility videos were taken from a parallel study (Balassy et al. 2018). Actin was purified from chicken pectoralis acetone powder (Pardee and Spudich 1982). Skeletal muscle myosin protein was purified from chicken pectoralis muscle (Sobieszek 1994) and phasic smooth muscle myosin was purified from chicken gizzard (Sobieszek 1994). Chicken gizzard and pectoralis were obtained from a local slaughterhouse. Myosin light chain phosphatase (MLCP) and myosin light chain kinase (MLCK) were purified from turkey gizzard (Sobieszek, Borkowski, and Babiychuk 1997) (gift from A. Sobieszek). Phasic smooth muscle myosin was phosphorylated prior to the motility study for 20 min at room temperature using 5.02mg/ml myosin, 4mM CaCl_2_, 12.5 mM MgCl_2_, 14.7 mM imidazole, 0.59 mM EGTA, 18mM DTT, 3.75 μM calmodulin, 5mM ATP and 0.07 μM MLCK. KCl and glycerol were added to final concentrations of 0.3M and 50% respectively. Actin was labelled with tetramethylrhodamine isothiocyanate-phalloidin (Sigma #P1951). Myosin, actin, and motility buffers are described in supplemental methods.

We used an inverted microscope (Eclipse Ti, Nikon) to observe the labelled actin with a 100x magnification oil immersion objective (Nikon CFI Plan Fluor 100X Oil), and recorded with an EM-CCD camera (Hitachi KP-E500) at 30 frames s^-1^ and a Matrox frame grabber (Matrox Mor 2VD-84) at an effective resolution of 0.124 μm/pixel. An X-Cite 120Q excitation light source was used. To maintain a temperature of 30 □ C in the flow through chamber an objective heater (Bioptechs) and heating plate slide holder (Chamlide TC) were used. Flow-through chambers consisted of a nitrocellulose coated coverslip with two strips of double-sided scotch tape on which a plastic sheet was positioned (Artus Corp, 0.125mm plastic shim) with a 2mm diameter small hole in the centre. A polycarbonate membrane (0.8μm pore size, Millipore Sigma #ATTP02500) was UV glued to the plastic, covering the hole. Injection of ATP or MLCP (3.5 μM) dissolved in motility buffer onto the membrane resulted in convection free diffusion into the flow-through chamber.

Motility assays were performed as described in (Hilbert, Cumarasamy, et al. 2013).

### Algorithm

Apart from parameter settings, the entire algorithm described below does not require any user input. For a wide range of motility videos recorded under different conditions, on two different set-ups and with a 40-fold difference in filament velocity and motile fraction we were able to use a single set of parameter settings. The software is written in Matlab (The MathWorks, Natick, MA) and published under a GNU General Public Licence v3, at https://github.com/Gijpma/Motility-of-Filament-ends

### Image optimization

Camera noise is reduced by averaging consecutive video frames. For all data shown here we found that averaging of 2 frames was sufficient. A 2D Gaussian smoothing filter with a standard deviation of 1 pixel is applied to the resulting image to further smooth out camera noise in the filament images; this allows for more accurate binarization (conversion to black and white). An image with a much stronger 2D gaussian smoothing (standard deviation of 20) is used as an indication of regional illumination and background noise variation. To correct for this uneven illumination, we subtract the heavily blurred image from the filament image, after which the resulting pixel intensities are scaled to fully use the dynamic range available in an 8-bit greyscale image. The image is then binarized by setting to a value of 1 (i.e. white) all pixels above an intensity threshold. The optimal threshold for detecting filaments is determined by analyzing the number of isolated white areas identified in the first frame for all possible threshold values (see Fig. 1A). For very low threshold values the entire image is connected, resulting in a single identified connected component, while gradually increasing the threshold value rapidly increases the number of identified components as noise pixels are isolated until a maximum is achieved (P1 in Fig. 1A). A further increase in threshold value rapidly decreases the number of detected noise pixels and thus connected components, until only the filaments remain. After this point the number of components either initially increases as the filaments break up into individual islands followed by a decrease to zero, or no initial increase occurs as the break up of filaments is balanced by the disappearance of filaments altogether. Because a clear local minimum filament count, corresponding to only filaments without noise, could not always be detected, we identify the first threshold value beyond P1 where the number of components drops below a parameter set value (here 300, labelled P2 in fig. 1A), and setting the threshold P3 at a parameter set value (here 5%) higher than P2. This method worked consistently in detecting filaments without spurious noise issues for over 300 videos recorded on two different motility set-ups and variable lighting conditions and motility quality without changing either of the two parameters.

**Fig 1:**
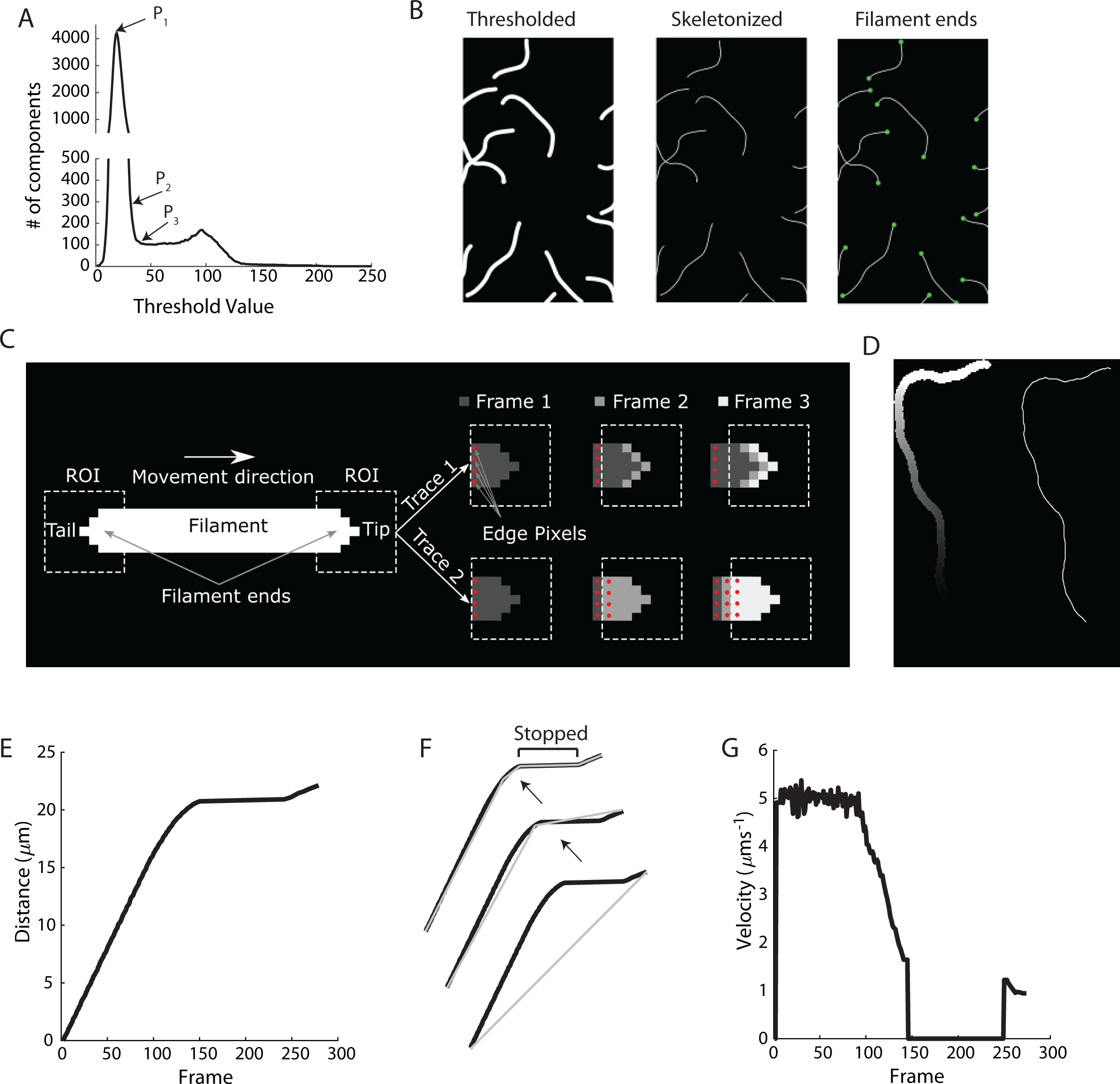
Motility of Filament Ends (MFE) algorithm details. A) Sample curve showing the number of components detected for all threshold values. P_1_ indicates maximum number of elements detected, P_2_ indicates point at which <300 elements are detected and P_3_ indicates the calculated threshold. B) Left side shows black and white image after thresholding (“thresholded”) with threshold value derived from A. Middle image shows the same image morphologically thinned (“skeletonized”). The right image shows filament end identification, indicated with green markers at filament ends (“Filament ends”). C) Filament movement trace generation. Left: thresholded image of a filament with identified filament ends, tip and tail, and the used Region of Interest (ROI) around the ends. Two traces are generated for each filament end. In frame 1 both traces store the whole ROI image, with the pixels at the edge of the ROI labeled as edge pixels (red dots). In frame 2, trace 1 only stores the pixels previously not occupied by the filament end, while in trace 2 all pixels of frame 2 are stored, overwriting some of the pixels from frame 1. This is repeated in frame 3, resulting in trace 1 showing a progression of the filament end, while trace 2 shows a progression of edge pixels. In this case, trace 1 is selected, by comparing the number of edge pixels in the final trace and used for further analysis. D) Sample of a single filament end trace (left) and a morphologically thinned and smoothed effective centreline (right). E) Frame vs. Distance calculated from filament end trace. A steep slope indicates fast movement, while a horizontal line segment indicates a stopped filament. F) Ramer-Douglas-Peucker algorithm for segmentation of the Frame-Distance trace. This algorithm segments the trace by starting as a linear connection between the start and end points and repeatedly breaking this line into sections connecting the most distant points on the curve from the line in the previous iteration. G) Resulting velocity vs. frame trace.

### Endpoint identification

After taking a threshold image (Fig. 1B Thresholded), all connected components (filaments) are skeletonized by repeated morphological thinning, i.e. removal of the perimeter pixels until a single pixel width line remains (Fig. 1B (Skeletonized). The endpoints and branchpoints of this skeleton image are identified (Fig. 1B Filament ends). Small branches are pruned, while big branches, indicating filament overlap, are kept. When an endpoint is in close proximity to a branchpoint, the endpoint is removed. This occurs when two filaments are overlapping, but the tip of one of the filaments is close to the body of the other filament. These points need to be removed to avoid analysis errors. Two filament ends in close proximity that do not belong to the same filament are also removed, while two filament ends in close proximity belonging to a single filament are analyzed as a single point by taking the average location of both endpoints. To track the movement of endpoints from frame to frame, the endpoints are cross-correlated between frames based on proximity only.

### Frame-to-Frame analysis

Consider the thresholded, idealized, and straight filament moving to the right as in figure 1C. In each frame, square Regions Of Interest (ROI) around the filament ends, large enough to capture the entire filament ends and a small part of the filament body, are stored (here 10×10 pixels) and the white pixels are time-stamped with the current frame number. To identify the direction of movement, the white pixels at the edge of the ROI are labelled as edge pixels (red dots in Fig. 1C). After all frames are read, two movement traces for each filament end are generated from the stored ROI images. Trace 1 stores for each frame, only the pixels, with their associated frame numbers, not previously occupied by the filament end. Trace 2 stores for each frame, all the edge pixels, with their associated frame numbers, thus overwriting all pixels that were previously occupied by the filament end. For movement to the right as in figure 1C, trace 1 shows a progression of the filament end, while trace 2 shows progression of the edge of the ROI. As we are interested in the progression of the filament end, only trace 1 is stored for this filament end. The other filament end, the filament tail, will show progression of the filament end with trace 2 and thus trace 2 is stored for this filament end. Identification of the right trace is done by comparing the number of edge pixels remaining in both traces.

Occasionally, a section of a trace shows a change in movement direction (i.e. a tip turns into a tail), which indicates “end-overtake”: two filament ends from different filaments move almost exactly over each other in opposite direction, confusing the algorithm into identifying this as a single end movement. These traces are identified by non-uniform distribution of the number of edge pixels and are rejected.

### Trace analysis

The trace (Fig. 1D, left) is reduced to a skeletal representation (single pixel width curve) by repeated morphological thinning (Fig. 1D, right). The resulting skeleton of the trace is pruned to remove any branches caused by off-track motion, image noise or a variation in lighting and focus (for details see supplementary methods). The resulting 2D curve is smoothed (a moving average filter on the x and y coordinates) to subpixel coordinates. This is necessary as the total path length is exaggerated by the restriction to whole pixel movement. Intensity values of the sub-pixel 2D curve coordinates are subsequently calculated by interpolation.

To derive a curve of frame number vs. distance of the filament end, we calculated the cumulative distance over the trace starting at the end with the lowest frame number (Fig. 1E). As the first frame endpoint image remains intact within the full trace (Fig. 1C, frame 3 Top), the frame number vs. distance curve will start with a distance increase of several pixels before an increase in frame number occurs. To remove this start-up effect (and end effect for tail traces), any points with multiple distance values for the same frame number are removed from the curve.

### Curve segmentation

Because of the coarseness of the distance vs. frame number curve, particularly for low velocity motility (<.1 pixel per frame), smoothing of the data is needed before velocities are calculated. However, motility assays often show stop-and-go motion, believed to be caused by dead motor heads, operation close to the cooperativity threshold of myosins with actin (Hilbert, Bates, et al. 2013) or non-permanent cross-linking with other proteins. Smoothing blends stopped and moving segments of the curve, creating artificially low moving speeds and shortened stopped sections. Here we first segment the curve into stopped and moving regions and apply smoothing only on the moving segments of the curve. The segmentation is done using the Ramer-Douglas-Peucker algorithm, which reduces the curve to segments in which none of the points on the original curve deviates further from the segment lines than a given error value (Fig.1F). For each segment, the velocity is calculated as the slope of the segment line, and any segment with a velocity below a given threshold value is set to stopped. Adjacent moving sections are merged, as well as adjacent stopped sections, after which the moving segments are smoothed with a moving average filter (Fig.1G). An average velocity of all moving filaments is calculated as well as the instantaneous motile fraction, i.e. the proportion of moving filaments at any point in time. To inspect analysis results a video is generated containing the original frames with green bars placed at each endpoint indicating the velocity of movement and red dots where endpoints are identified as stopped (for example, see Fig. 4C)

**Fig 2.**
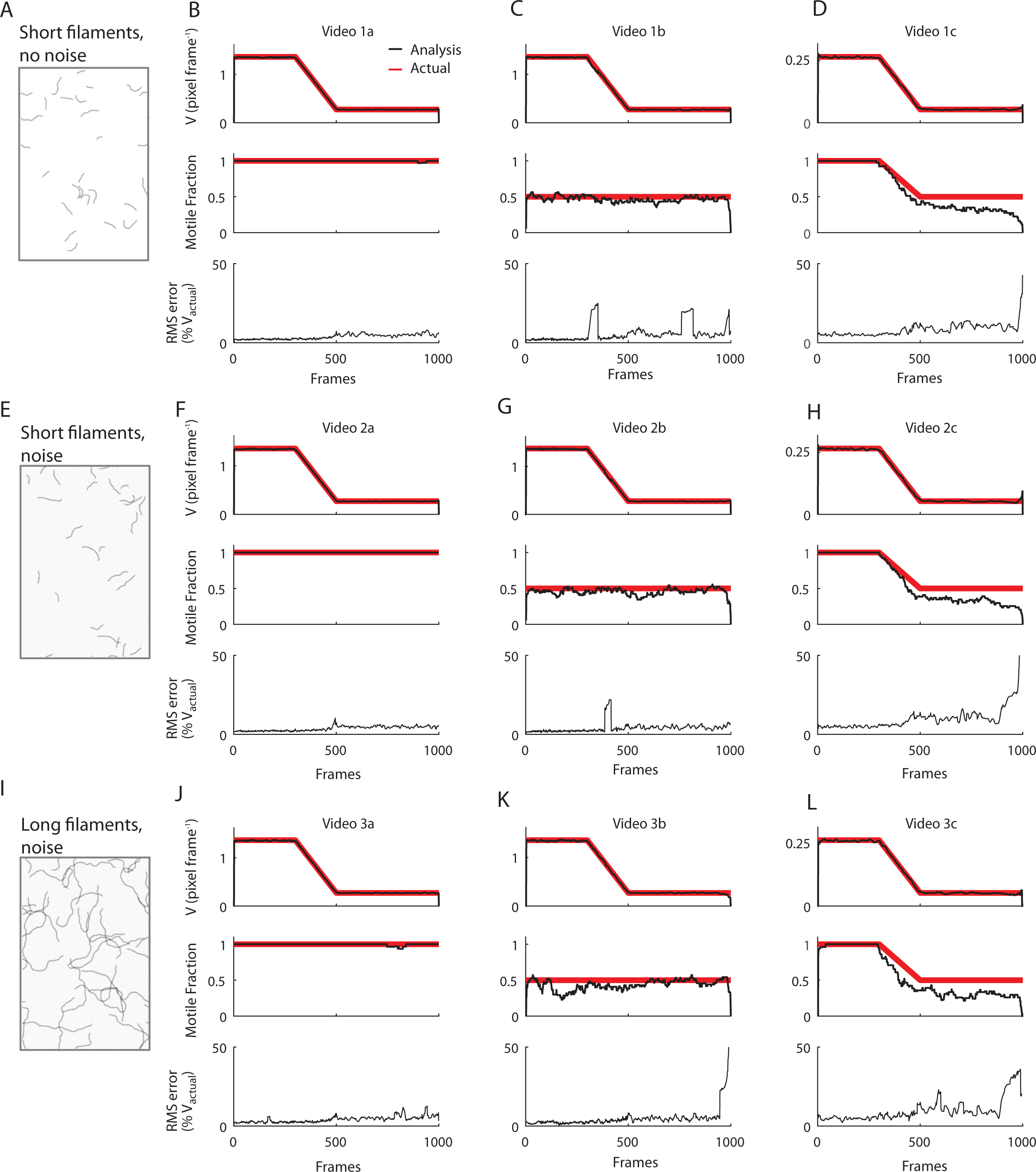
Testing of the MFE algorithm with several mock filament motility videos. A) inverted single frame of mock video used, containing curved filaments with no image noise (video 1a-c, see text). B), C) and D) show the estimated (black) and actual (red) velocity profile (top), measured and actual motile fraction (middle) and fractional RMS error of individual traces in the velocity (bottom) for three different velocity and motile fraction profiles as indicated by the video name (see text). E) inverted single frame of mock video with curved filaments as in A) but with image noise (gaussian speckle, video 2a-c). F), G), and H) velocity, motile fraction and RMS error data for same velocity and motile fraction profiles as in B), C) and D). I) inverted single frame of mock video with long curved filaments with extensive filament overlap and noise (video 3a-c). J), K) and L) show velocity, motile fraction and RMS error as in B), C) and D). Image noise and extensive filament overlap do not affect RMS error of the individual traces nor the average velocity. RMS error is increased for very low velocities, and motile fraction is underestimated at these velocities.

**Fig 3:**
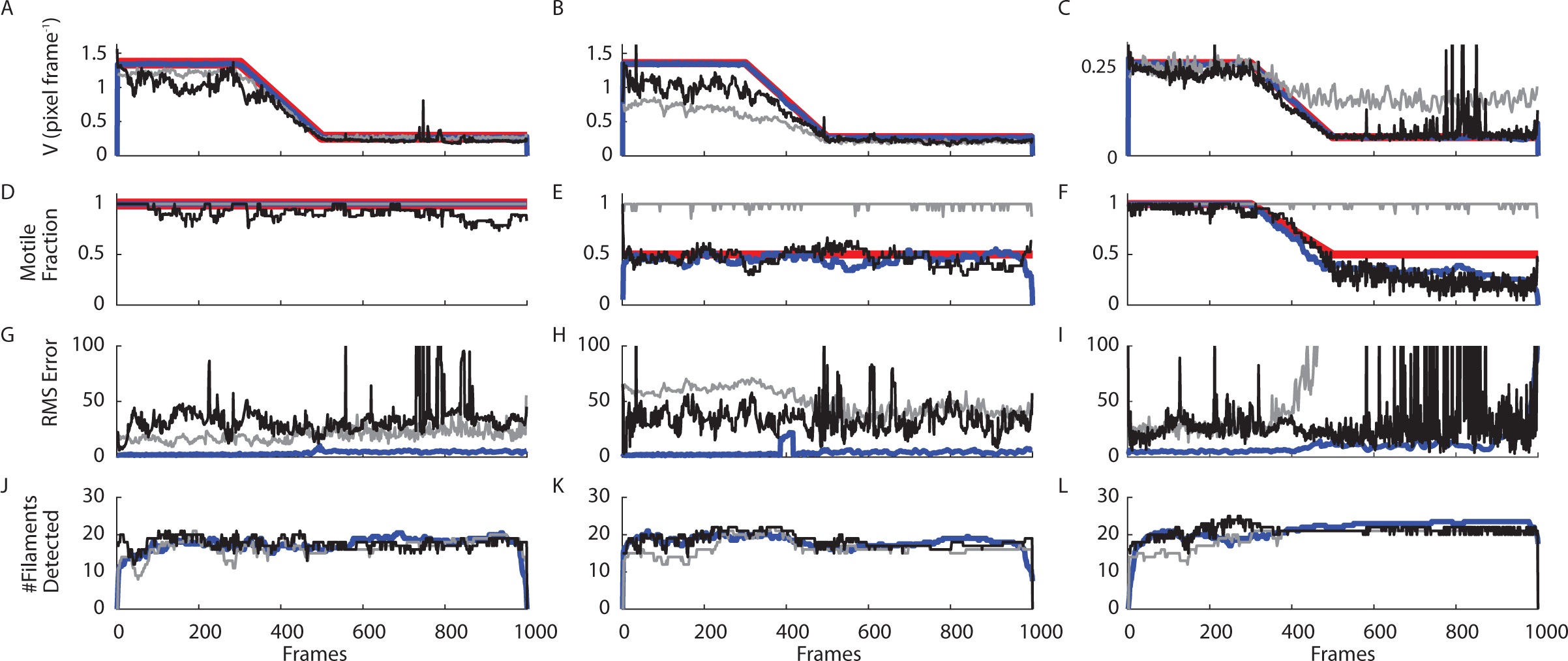
Comparison with Fiesta and Centroid tracking algorithms for mock videos of figure 2 F-H. Fiesta and Centroids tracking algorithm used manual trace dropping of erroneous traces (not correctly identified crossing traces or picked up image noise). A), B) and C) Velocity traces of the three algorithms and the actual velocity of the mock videos used. D), E) and F) Motile fraction for the three algorithms and actual motile fraction. Centroids tracking overestimates motile fraction using the 0.04 pixel frame^-1^ cut-off velocity used for all three algorithms. Fiesta and MFE both measure motile fraction accurately except for very low velocities. G), H), and I) fractional RMS error of the individual filament traces. Error is consistently much lower for the MFE algorithm. Note that the centroids algorithm gives a minimum velocity of ∼0.18 pixel frame^-1^ as a result of image noise, and thus the RMS error at an actual velocity of 0.05 pixel frame^-1^ (0.2 μm s^-1^) in figure 3I is well over 100%. J), K), and L) number of filaments detected in the three videos. Each video contained 25 filaments, including filaments at frame edges and crossing filaments. For the MFE algorithm the number of filament ends was divided by 2 to get to the number of filaments.

**Fig 4.**
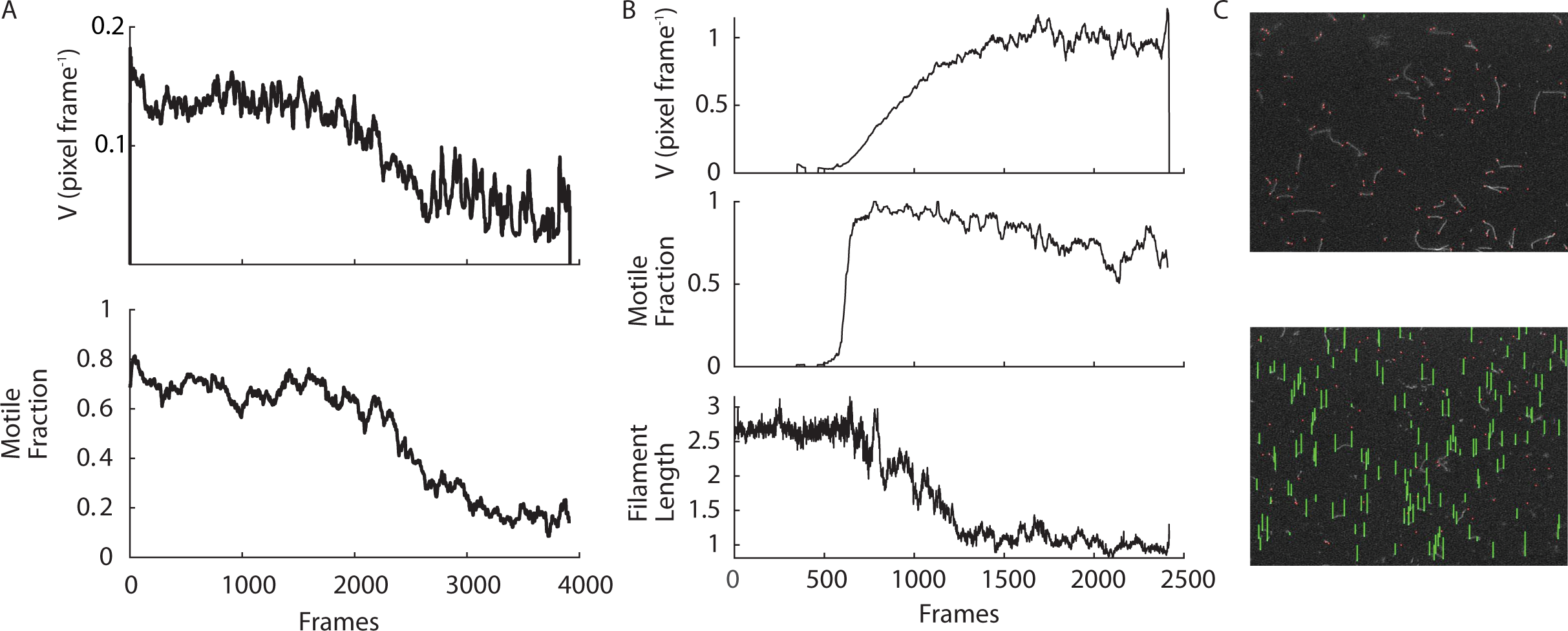
MFE performance on actual motility videos. A) Velocity (top) and motile fraction (bottom) profile of actin on phasic myosin in which myosin light chain phosphatase (MLCP) diffuses into the chamber starting at frame 0. Video resolution is 0.124 pixel μm^-1^ at 30 frames s^-1^ B) Velocity (top), motile fraction (middle) and length (bottom) profile of actin motility on skeletal muscle myosin in which ATP starts to diffuse into motility chamber at frame 0. C) 2 frames from output video of MFE algorithm of video from B) showing red markers on stopped filament ends and green bars indicating velocity of moving filament ends. Top is frame 150, bottom is frame 2000.

### Mock motility video generation

We developed a Matlab algorithm for the creation of mock motility videos with known velocity and motile fraction profiles over time, allowing for the evaluation of the accuracy of the proposed method and comparison with existing methods in the literature. To accurately represent filament motility videos, the video generation software allows for control of velocity, motile fraction (including stop-and-go movement), variable distribution of filament size and curvature as well as various modes of background noise. For a detailed description of the methods used, see supplementary methods. For sample videos, see supplemental videos “3C” and “4C”.

### Mock videos

We tested several mock videos, gradually increasing the level of complexity (see Fig 2 and supplementary materials, 30 frames s^-1^ and 0.124 μm pixel^-1^):

1. Curved filaments; no noise;
2. Curved filaments; noise;
3. Long, overlapping curved filaments; noise

For each of these 3 scenarios, 3 velocity regimes were created:

a. Variable velocity (changing from 5 to 1 μm s^-1^)
b. Variable velocity with stop-and-go movement (velocity as in a, motile fraction 0.5).
c. Low variable velocity with variable motile fraction (velocity changing from 1 to 0.2 μm s^-1^, motile fraction as in b

The generated videos were sampled at 30 frames s^-1^, with 25 generated filaments in each frame and changes in velocity and motile fraction were applied between 300 and 500 frames (see figure 2). We calculated root mean squared (RMS) error from the measured difference in velocity of each filament relative to the input velocity of the mock motility video at each frame.

## Results/Discussion

### Mock Video Data

We tested the performance of the proposed Motility of Filament Ends (MFE) algorithm on 9 mock motility videos of 1000 frames each. On a 4-core processor using parallel processing of the videos this took 205s. Figures 2A-D show that the RMS noise of the individual filament traces is slightly increased for low (Fig. 2B) and very low velocities (Fig. 2D). Note that the RMS error measures individual filament error, while the velocity values in figure 2 are an average of all detected filaments in the video. Consequently, a larger RMS error does not necessarily correspond to a greater error in the average velocity reading shown here. The introduction of stop-and-go movement does not increase the noise (Fig. 2C). At very low velocities the motile fraction is underestimated towards the end of the video. Introduction of noise only slightly increases the noise of the analysis as shown in figures 2F-H. Increasing filament length to the point of extensive filament overlap did not decrease the accuracy of the analysis substantially (Fig 2I-L).

The mock video tests show that the algorithm is capable of accurately detecting a wide range of velocities down to movements less than 0.05 pixel frame^-1^ (0.2 μm s^-1^, 0.124 μm per pixel). With a cut-off velocity for moving vs. stopped filaments set at 0.04 pixel frame^-1^, some filament ends that are detected only towards the end of the video can be mistakenly identified as stopped, resulting in the drop off in motile fraction in Figs. 2D, H and L. If stop-and-go motion occurs with relatively short moving or stopping times the analysis may identify the motile state of filament ends incorrectly as well. Most variability in individual filament end velocity occurs towards the start and end of the filament end’s trace as the tip and tail of a skeletonized trace are most sensitive to noise.

### Comparison with existing algorithms

We subsequently compared the performance of the MFE algorithm with two existing algorithms on either side of the complexity spectrum: a filament centroid tracking algorithm previously developed at our lab (Hilbert, Bates, et al. 2013) and a filament fitting algorithm (FIESTA, (Ruhnow, Zwicker, and Diez 2011)). We previously developed an algorithm based on the motion of filament centroids, which can do both manual trace dropping for overlapping filaments, or an automated procedure based on machine learning (Hilbert, Cumarasamy, et al. 2013; Hilbert, Bates, et al. 2013). Here we used its manual trace dropping feature for the cleanest results. The FIESTA algorithm aims to find filament locations at subpixel accuracy by dividing detected filaments into small sections containing a filament tip, a filament body or a crossing filament and then fitting models of these features (Ruhnow, Zwicker, and Diez 2011). For a specific set of conditions, the authors reported location tracking accuracies on the order of 10 nm. The FIESTA software has the benefit of being able to handle crossing filaments as well as the subpixel accuracy, at the cost of a much greater computation time. However, in tracking the velocity of actual moving filaments the authors showed velocity variability of ∼250 nm/s. As initial results showed very large position variability in a subset of filaments in the FIESTA software, likely because of incorrectly identified filaments, we also applied here a manual trace dropping to clean the data. Run time for 3 videos of 1000 frames each was 1650s. Any trace dropping in the proposed MFE algorithm was fully automated and based on criteria specified in the methods. To ensure equal comparisons, the same cut-off velocity to distinguish between stopped and moving filaments (0.15 μm s^-1^, 0.04 pixel frame^-1^ in the MFE algorithm was used for the FIESTA and centroids tracking software as well.

Neither the FIESTA nor the Centroids tracking algorithm were able to detect movement in the long crossing filament videos used for figure 2 I-L. Figure 3 shows that for mock videos of curved filaments with noise for the same three velocity and motile fraction scenarios as in figure 2 E-H, the RMS error for the MFE algorithm is considerably less than both the FIESTA and centroids tracking algorithms for all three videos. Also, the error in both the centroids and FIESTA algorithms results in a reduction in the measured average velocity. Note that we introduced a 20-frame moving average smoothing to the output of the Centroids and Fiesta algorithms to allow a fair comparison with the MFE algorithm which has a similar smoothing built in. The data also show that the introduction of stop-and-go movement greatly decreases the accuracy of the velocity signal of both the FIESTA and the centroid tracking algorithms, as stopped filaments can have an occasional apparent velocity above the set cut-off value because of the video noise. A similar issue arises for low velocities (Fig. 3 C, F and I) where particularly the FIESTA algorithm results in unstable velocity estimations. At very low velocities the MFE algorithm estimates motile fraction much more accurately then both the centroids and FIESTA algorithms. The FIESTA algorithm does show an improved accuracy of the motile fraction estimation at the end of the video during slow motility as the distinction of stopped vs moving filaments is made instantaneously rather than over a timespan requiring a minimal movement of one pixel.

### Application to actin motility data

We tested the MFE algorithm on 2 sets of actual data with changing velocity and motile fraction profiles. These videos are part of a future publication for which the current algorithm was developed. In the first video we injected MLCP into a motility assay of phasic smooth muscle myosin propelling actin filaments. As this myosin is relatively slow (0.5 μm s^-1^ starting velocity, 0.13 pixel frame^-1^ at 30 frames/s) the cut-off velocity was set lower to 0.03 μm s^-1^ (0.008 pixel frame^-1^) to ensure accurate detection of any decreases in velocity as is expected to occur in response to dephosphorylation of the regulatory light chain of the myosin molecules. The video contained sudden focus and illumination changes occurring around the time of injection of MLCP, which did not affect the filament resolution by the algorithm. The MFE software shows a gradual decrease in motile fraction in parallel with a gradual decrease in velocity, with good filament resolution (Fig. 4A). The very low velocities combined with stop-and-go motion are unlikely to be detected by existing algorithms based on frame-to-frame or trace analysis. In a second video we injected ATP into an actin motility assay with skeletal myosin. As the actin is originally in a rigor state, the filaments have a motile fraction of zero and a velocity of zero. As the ATP enters the motility chamber through a diffusion membrane (see methods) the ATP concentration in the chamber gradually increases, resulting in increasing motile fraction and velocity of the actin filaments. Our data clearly show that motile fraction increases rapidly to a plateau level while velocity increases much more gradually, taking over 30 s to plateau (Fig. 4B top and middle). This indicates that cross-bridge cycling is rate-limited by the amount of available ATP, but that the diffusion of ATP is uniform across the field of view. The video shows rapid break-up of filaments, which may be caused by the gradual activation of the myosin heads as the concentration of ATP increases. This would result in active heads pulling on actin while remaining rigor heads hold on, resulting in breakage of the filament. The algorithm detected the change in average length (Fig. 4B bottom).

### Limitations

While the proposed algorithm performs well in most situations, under certain conditions filament movement could be misidentified, or filament ends are not properly detected.

- During stop-and-go movement the ability of the algorithm to detect stopped periods is dependent on the cut-off velocity, the image resolution and video noise. As a result, average velocities may be underestimated during rapid stop-and-go turnover.
- The automated threshold detection algorithm uses only the first frame, with a correction for overall illumination changes throughout the video. If the first frame is not representative of the remainder of the video (primarily if the focus is different) the algorithm will have difficulties.
- While non-directional motion is mostly filtered out by the trace detection approach, slow, random motion due to unattached filament ends may be detected as directional motion, particularly for short traces.

## Conclusions

We showed that the MFE algorithm is capable of accurate tracking of variable velocity motility assays with a high time resolution and superior accuracy to two of the leading algorithms for tracking filament motility. It does this robustly with minimal complexity, user input and CPU time. The focus on filament ends instead of entire filaments also allows for the unique ability to detect filament motion in assays with long, overlapping filaments. The history retaining method of filament end trace detection combines the benefits of trace-based analyses (detection of directed motion, low velocities detection) with the benefits of frame-to-frame filament location analyses (time resolution).

## Supporting information

see supplemental videos ???????3C??????? and ???????4C???????

see supplemental videos ???????3C??????? and ???????4C???????

